# QTLs analysis for cold tolerance in rice (*Oryza sp*.) at the seedling stage using F_2_ population developed from cross between the genotypes NERICA L-19 and Diamante

**DOI:** 10.1101/2022.03.15.484433

**Authors:** Branly Wilfried Effa Effa, Baboucarr Manneh

## Abstract

Rice (*Oryza sp*.) is the most important food crop of the developing world and the staple food of more than half of the world population. It is produced in about 110 countries, including Africa under traditional production systems where field conditions are often variable and yield is low. This is due to the preponderance of abiotic stresses such as cold in these systems. The aim of this study was to identify QTLs associated with cold tolerance at the seedling stage. It was obtained: a phylogenetic tree showing individuals tolerant and sensitive to cold in the Sahel conditions at the seedling stage; 5 majors QTLs associated with cold tolerance located on chromosome 4; QTLs associated with cold tolerance at the seedling stage using F_2_ population developed from cross between NERICA L-19 X Diamante. All these results are interesting, because to pass local varieties sensitive to cold tolerant varieties through the identification of QTLs associated with cold tolerance that might be introduced. This will enable African producers; especially small producers which have low resource to buy a plant material adapted to the cold period to optimize their rice yields.

## INTRODUCTION

Rice, the major cereal crop cultivated worldwide [1] can be classified into two major cultivar types from *Oryza sativa* species: indica and japonica. Indica cultivars are grown mostly in hot and humid tropical low lands; in contrast japonica cultivars are grown in temperate and sub-temperate regions and high altitude areas of tropics [2]. There is also the existence of an African species: *Oryza glaberrima* [3]. Two species of rice have been crossed, producing a promising hybrid “NERICA” [4].

Rice is a cold sensitive plant, low temperature effect is manifested at different growth stages such as germination, seedling, vegetative, reproductive, and grain maturity [5-8]. Rice plants are injured at seedling stage when they are grown below 25°C [9], however in West Africa average annual minimum temperatures are between 16 °C and 20 °C [10]. Screening for cold sensitivity of different rice genotypes in breeding programs commonly relies on visual observations under natural field conditions [5, 11]. Methods were also developed to assess cold tolerance under controlled conditions [5]. These include chlorophyll chilling survival tests using 10°C at the two leaf stage [12], measurement of radicle growth [13] or seedling vigor [14]. Previous studies have indicated that rice response to low temperature stress is complex and certainly controlled by more than one gene [5]. Objective of this study was to identify QTLs associated with cold tolerance at seedling stage. We used linked Simple Sequence Repeat (SSR) marker analysis in two batches of rice material, firstly 30 varieties for allelic diversity study, and secondly 180 F_2_ individuals derived from cross between a cold tolerant Chilean cultivar DIAMANTE [15] and a Cold susceptible African cultivar NERICA L 19 to identify QTLs associated with cold tolerance at seedling stage.

## MATERIALS AND METHODS

### A. plant materials

This study was led to Sahel regional station of AfricaRice located at Saint Louis (Senegal). Two batches of rice materials were used in this study first consisted of 30 varieties phenotypically classified as sensitive and cold tolerant varieties. These genotypes were used to conduct an allelic diversity study for five major QTLs associated with cold tolerance in rice at the seedling stage.

Second batch consisted of 180 F2 populations from cross between a cold sensitive African cultivar NERICA L-19 and a cold tolerant Chilean Diamante, and their two parents present on each block. The design consisted of 10 incomplete blocks and each block contained 20 individuals. This batch was used to identify Diamante QTLs involved in cold tolerance in the background of an African variety “NERICA L 19”.

### B. evaluation of cold tolerance

F_2_ were transferred 11 days after transplanting at 2-3 leaf stage in a cold room for screening. Treatment in the cold lasted seven days. Treatment was stopped when the susceptible control (NERICA L 19) had reached a level of wilting which earned him a score of 9 in the Standard Evaluation System (SES) for rice that was developed by the International Rice Research Institute [16]. Rate of plant growth before and after stress was estimated. F_2_ were sown in the experimental Augmented Design [17]. Behavior of individuals in trays (F_2_ and both parents present on each block) was analyzed.

### C. statistics analysis

Score of cold effect on seedling rice according to SES code [16], plant height and leaf length to each individual. The biometrics parameter values were adjusted according to the formula [adapted from 17]:

**r**_**j**_ **= B**_**j**_**-M**, where r_j_ = Effect of j^th^ block; B_j_ = Average of all witnesses in the j^th^ block; M = Medium Large witnesses, **Adjusted number = (Number unadjusted) – r**_**j**_

ANOVA were performed with SAS 9.2 software using the GLM model (Proc Insight) and the software GenStat Discovery Edition 4 by applying the model Mixed Models (Linear Mixed Models) [18]. Heritability of plant height, and leaf longest length were calculated before the screening and after the screening with GenStat; setting ‘‘genotype’’ and randomizing ‘‘block’’. The formula used for these calculations was:

**Heritability= [Vg / (Vg + Ve)] X 100**; With, Vg= Variance due to differences between genotypes; Ve = variance due to environmental effects caused by errors on the parameters.

### D. genotyping and molecular analysis

Leaves harvest was conducted in two steps: 25 days after transplanting in the greenhouse for each of the 30 genotypes used in the study of allelic diversity, and the 7^th^ day of screening in a cold room for the 180 F2 used for QTLs identification. Genomic DNA was extracted by a modified CTAB. Pure DNA was diluted before use to 20%. The unit mix used for PCR was Ultra-Pure Water: 4μl; 1.5 mM PCR buffer (10X Buffer): 1μl; 1 mM dNTPs: 1μl; 25 mM MgCl_2 0.5_ μl; Forward: 0.5 μl; Reverse: 0.5 μl; 3% DNA polymerase: 0.5 μl; 20% pure DNA: 2μl. Program used for PCR was Alamgir SSR 55, a 34 cycle’s program. 30 SSR markers were used for allelic diversity study. Analyzes were performed using Darwin5 software to draw a phylogenetic tree by the method of distance (NEIGHBOR-JOINING) for allelic diversity study. Then to identify the most discriminative markers, gene diversity index were calculated according to the method of [19] using the following formula:

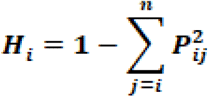

P_ij_ is the frequency of the j^th^ allele for the i^th^ marker and n is the number of alleles observed in the study population.

For QTLs analysis in F2 population, 159 SSR markers were used.

### E. qtls identification

QTL analysis was performed with qGene 4.3.10 software using Simple Interval Mapping (SIM) option and Composite Interval Mapping (CIM) option to validate presence of putative QTL. To justify presence of a putative QTL, LOD ≥ 2 was considered [20].

## RESULTS

### A. Evaluation of cold tolerance

Analysis of F_2_ biometric parameters showed that data taken before screening and those taken after screening, height and length of the longest leaf have a normal distribution (Figures 1 and 2).

**Figure 1:**
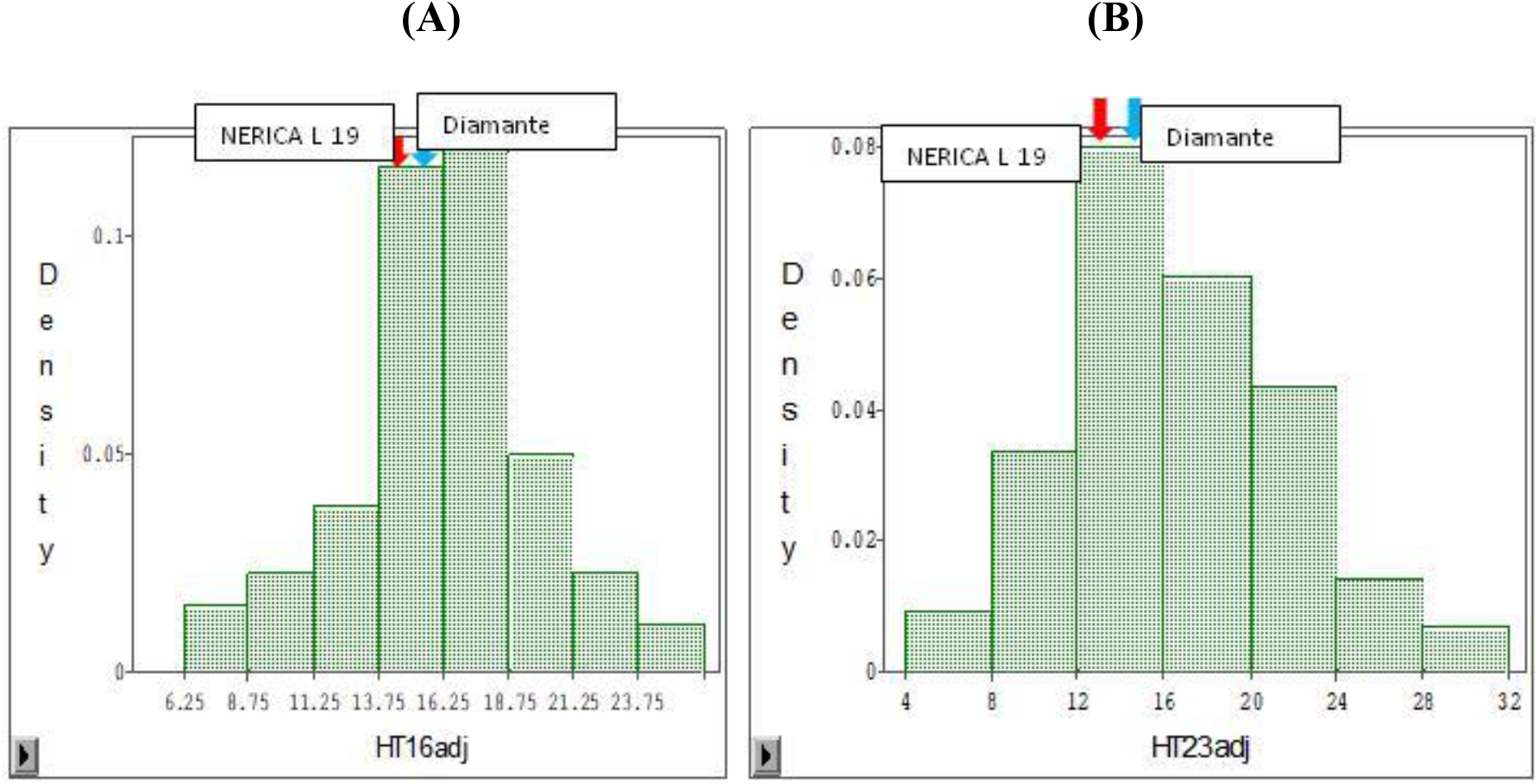
adjusted Height: (A) before screening: Average =16.33 cm, Sd =3.62, CV=22.15; (B) after screening: Average=16.56 cm, Sd=5.02, CV=30.33. Red arrow for Nerica L 19 genotype; blue arrow for Diamante genotype; Sd : Standard Deviation; CV : variation Coefficient.

**Figure 2:**
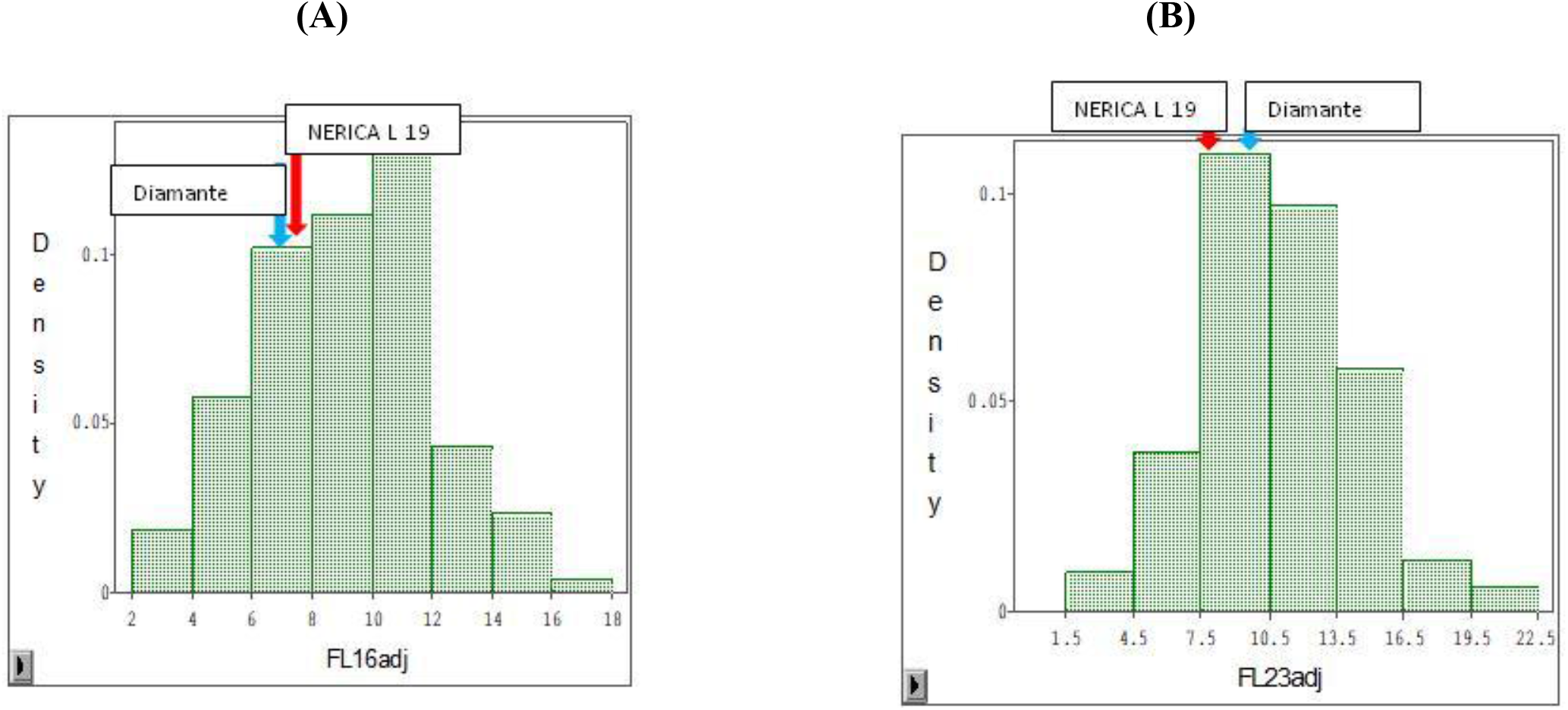
leaf length adjusted : (A) before screening: Average=9.17 cm, Sd=2.9, CV=31.67; (B) after screening: Average=10.92 cm, Sd=3.64, CV=33.37. Red arrow for Nerica L 19 genotype; blue arrow for Diamante genotype; Sd : Standard Deviation; CV : variation Coefficient.

While score has a distribution skewed to the susceptible control NERICA L 19 (Figure 3). Average heights before and after screening were respectively 16.33 cm and 16.56 cm. Average lengths of leaves before and after screening were respectively 9.17 cm and 10.92 cm. Average score was 6.87 at the end of screening, which has been likened to 7. NERICA L-19 and Diamante showed sensitive behavior with respective scores of 9 and 7, Diamante is less sensitive than NERICA L-19.

**Figure 3:**
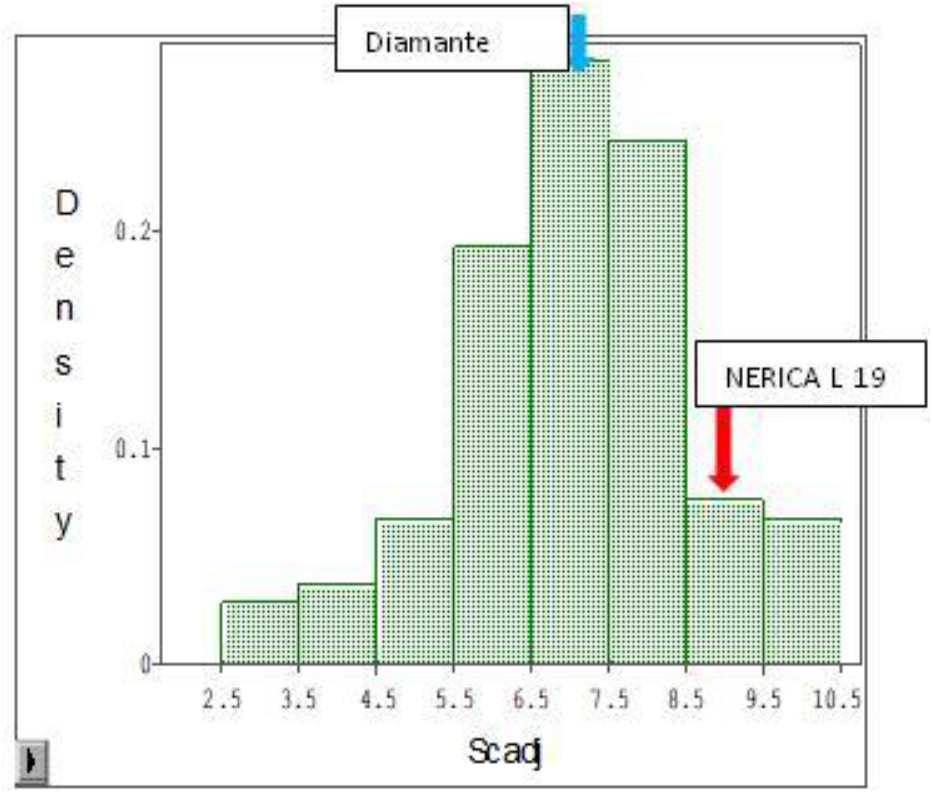
F_2_ population (derived from a cross between NERICA L-19 and Diamante) adjusted score Distribution after cold screening. Average=6.84 cm, Sd=1.57, CV=22,98. Red arrow for Nerica L 19 genotype; blue arrow for Diamante genotype; Sd : Standard Deviation; CV : variation Coefficient.

ANOVA (Table 1 and 2) was used to determine contribution of different treatments (genotypes and blocks) on measurement parameters variability.

**Table 1:**
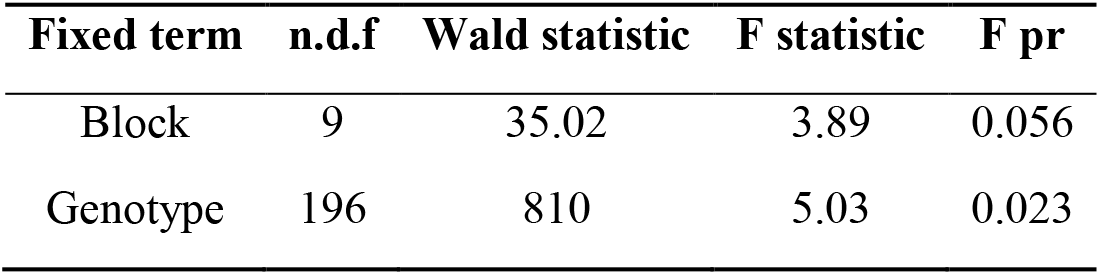
ANOVA for height after screening

**Table 2:** ANOVA for leaf length after screening

| Fixed term | n.d.f | Wald statistic | F statistic | F pr |
| --- | --- | --- | --- | --- |
| Block | 9 | 128.55 | 14.28 | 0.002 |
| Genotype | 196 | 2507.27 | 15.57 | 0.001 |

Then an analysis of heritability was performed to determine proportion of total variability of a parameter contributing to differentiate the F_2_ plants (effect of genotype). In terms of plant height and leaf length, heritability was higher before screening than after (Table 3).

**Table 3:**
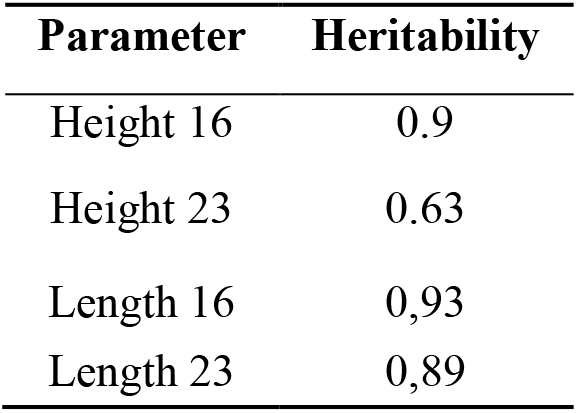
Heritability of plant height and leaf length of F2 population

In addition, correlation analysis allowed knowing relationship between plant height and leaf length before and after screening, between height and rate of growth in height after screening, and between leaf length and leaf rate growth after screening (Figure 4 and 5). Relationship between height and leaf length before and after screening was linear and highly significant (R = 0.82, P < 0.001 and R = 0.92, P < 0001 respectively). However, relationship between plant height and height growth, and relationship between leaf length and leaf length growth were quadratic and highly significant (P < 0001).

**Figure 4:**
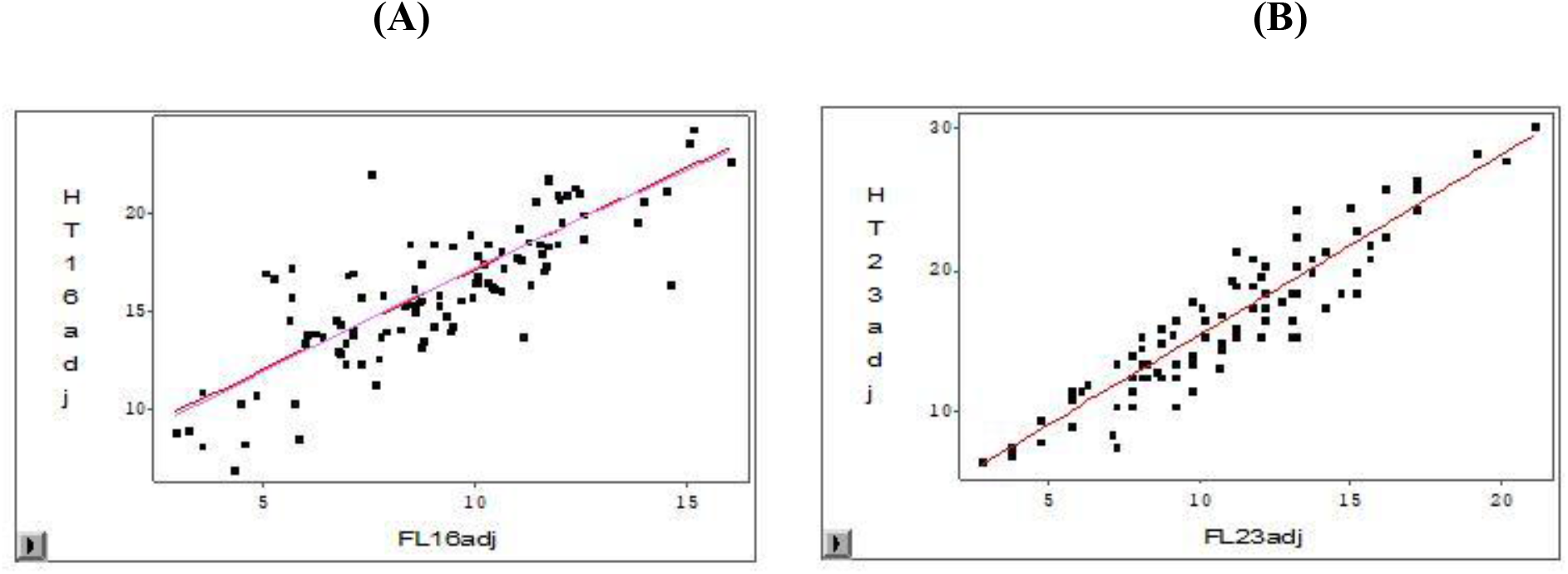
height function of leaf length: (A) before screening: Y=1,0259x + 6,9231; R=0,82; (B) after screening: Y=1,2676x + 2,7162; R=0,92.

**Figure 5:**
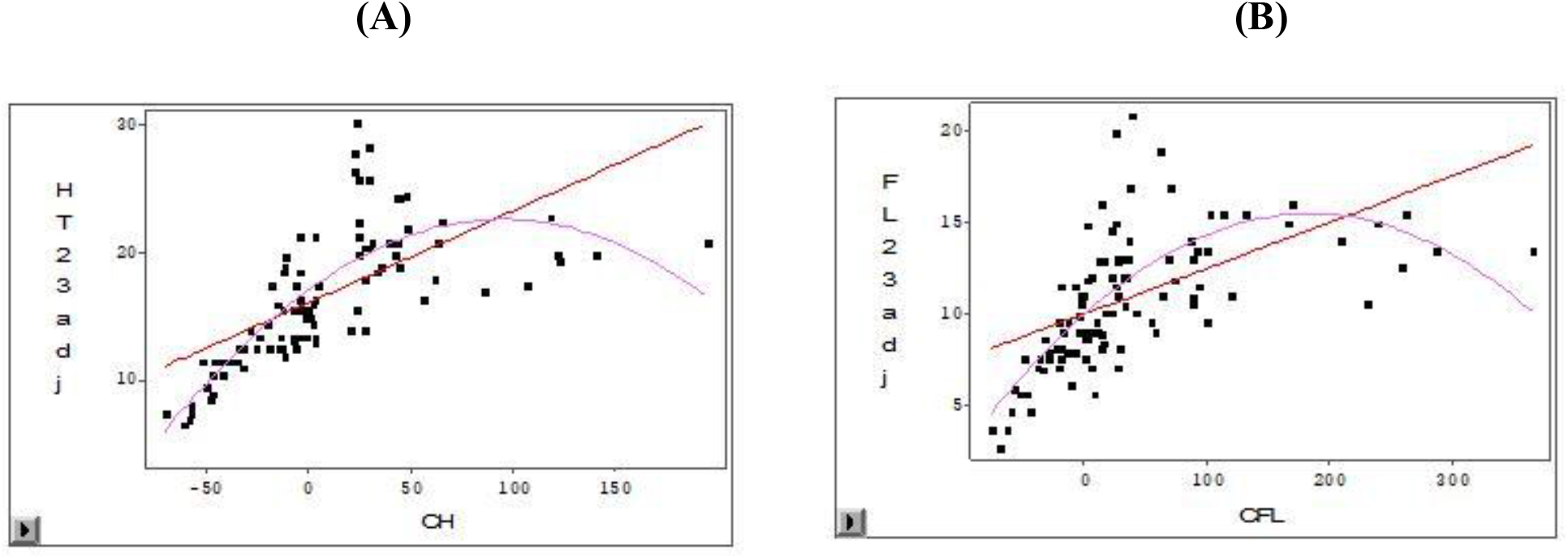
(A) height after screening function of height growth: Y = -0,0006x^2^ + 0,1183x +16,9255; R=0,79; (B) leaf length after screening function of leaf growth: Y = -0.0002x^2^ + 0.0601x + 9.9535; R=0,71

### B. Genotyping and molecular analysis

Index of genetic diversity of each marker used in this study was calculated, it varies from 0.37 to 0.86. The lowest (0.37) index was obtained with SSR marker RM101 and the highest (0.86) with RM335 SSR marker. Most discriminative were 13, each having a genetic diversity index greater than 0.72 (Table 4).

**Table 4:** Index of genetic diversity of discriminative markers

| Index of genetic diversity | Marker | Chromosome | Position (cM) |
| --- | --- | --- | --- |
| 0.86 | RM335 | 4 | 21.5 |
| 0.84 | RM348 | 4 | 137.9 |
| 0.82 | RM261 | 4 | 35.4 |
| 0.81 | RM241 | 4 | 106.2 |
| 0.8 | RM255 | 4 | 135.4 |
| 0.78 | RM19 | 12 | 20.9 |
| 0.77 | RM427 | 7 | 1.1 |
| 0.76 | RM292 | 1 | 66.4 |
| 0.76 | RM436 | 7 | 0 |
| 0.75 | RM230 | 8 | 112.2 |
| 0.74 | RM231 | 3 | 11.5 |
| 0.73 | RM270 | 12 | 91.3 |
| 0.73 | RM296 | 9 | 0 |

Analysis of allelic diversity of 30 varieties with SSR markers yielded a phylogenetic tree (Figure 6) which shows two major distinct groups. Above represent susceptible varieties cold and below tolerant varieties. In each group, there are several sub-groups according to their genetic proximity as revealed ramifications.

**Figure 6:**
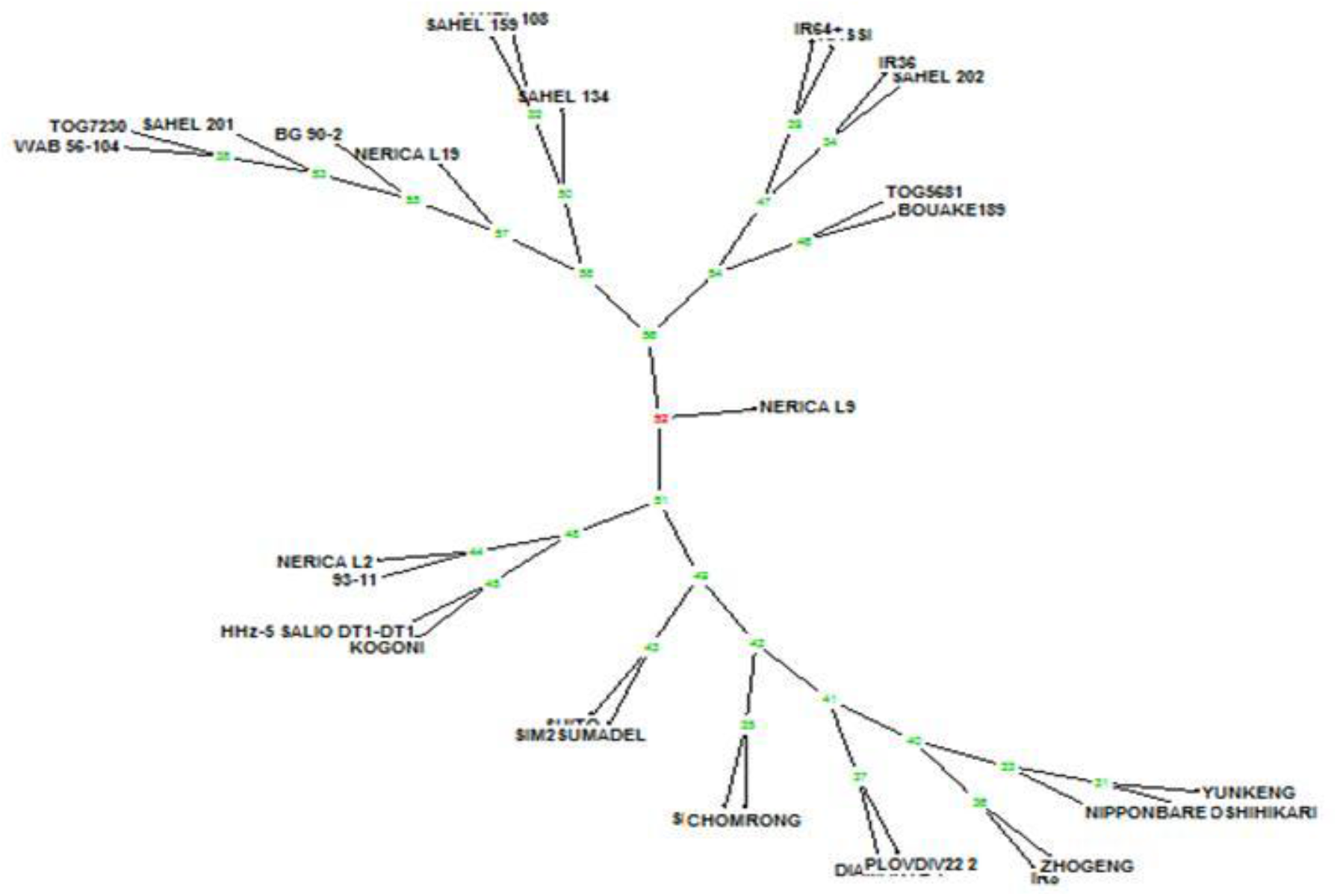
Phylogenetic tree for cold

### C. QTLs identification

Putative QTLs were identified using qGENE software independently of program performed, SIM or CIM (table 5 and table 6). Among the two programs, CIM program will detect more putative QTLs than SIM program. Thus, SIM program allow detecting 14 putative QTLs, 3 on chromosome 1; 2 on chromosomes 6, 10 and 12; and 1 chromosomes 3, 4, 7, 9 and 11. While CIM program has detected 15 putative QTLs, 3 on chromosomes 1 and 6; 2 on chromosomes 10 and 12; and 1 on chromosomes 3, 4, 7, 9 and 11.

**Table 5:** Putative QTLs obtained with SIM program

| Program | QTLs | chromosome | Position | marker | LOD | Parameter |
| --- | --- | --- | --- | --- | --- | --- |
| SIM | qCTS1a | 1 | 22.6 | RM220 | 2,5 | FL16adj |
|  | qCTS1b | 1 | 54.2 | RM250 | 3,5 | HT16adj,<br>HT23adj,<br>FL16adj,<br>FL23adj,<br>CFL, CH |
|  | qCTS1c | 1 | 115.2 | RM246 | 2 | FL16adj |
|  | qCTS3a | 3 | 215.5 | RM586 | 2 | HT16adj |
|  | qCTS4a | 4 | 106.2 | RM241 | 2,3 | FL23adj |
|  | qCTS6a | 6 | 13.5 | RM225 | 2,2 | CH |
|  | qCTS6b | 6 | 121.6 | RM528 | 3 | CH, CFL,<br>HT16adj,<br>HT23adj,<br>FL23adj |
|  | qCTS7a | 7 | 88.2 | RM234 | 2,2 | Scadj |
|  | qCTS9a | 9 | 99.4 | RM215 | 6,6 | HT16adj,<br>FL16adj,<br>CFL, CH,<br>HT23adj,<br>FL23adj |
|  | qCTS10a | 10 | 25.2 | RM239<br>(RM311) | 2 | Scadj |
|  | qCTS10b | 10 | 110.4 | RM333 | 2,5 | Scadj |
|  | qCTS11a | 11 | 50.8 | RM479 | 3 | Scadj |
|  | qCTS12a | 12 | 43.2 | RM512 | 2,3 | CH, CFL,<br>HT16adj,<br>HT23adj,<br>FL23adj |
|  | qCTS12b | 12 | 74.5 | RM309 | 3,3 | CH |

qCTS= Putative QTLs associated with cold tolerance in rice at the seedling stage, the numbers correspond to
chromosomes on which they were detected and letters order. HT16adj: height adjusted before screening; FL16:leaf
length adjusted before screening; HT23adj: height adjusted after screening; FL23adj: leaf length adjusted after
screening, CH: growth in height; CFL: leaf growth; Scadj: adjusted score.

**Table 6:** Putative QTLs obtained with CIM program

| Program | QTLs | chromosome | Position | marker | LOD | Parameter |
| --- | --- | --- | --- | --- | --- | --- |
| CIM | qCTS1a | 1 | 22.6 | RM220 | 2,5 | FL16adj |
|  | qCTS1b | 1 | 54.2 | RM250 | 3,6 | HT16adj,<br>HT23adj,<br>FL16adj,<br>FL23adj,<br>CFL, CH |
|  | qCTS1c | 1 | 115.2 | RM246 | 2 | FL16adj |
|  | qCTS3a | 3 | 215.5 | RM586 | 2,1 | HT16adj |
|  | qCTS4a | 4 | 106.2 | RM241 | 2,5 | FL23adj |
|  | qCTS6a | 6 | 13.5 | RM225 | 2,2 | CH |
|  | qCTS6c | 6 | 99 | RM454 | 2 | CH |
|  | qCTS6b | 6 | 121.6 | RM528 | 3 | CH, CFL,<br>HT16adj,<br>HT23adj,<br>FL23adj |
|  | qCTS7a | 7 | 88.2 | RM234 | 2,5 | Scadj |
|  | qCTS9a | 9 | 99.4 | RM215 | 6,6 | HT16adj,<br>FL16adj,<br>CFL, CH,<br>HT23adj,<br>FL23adj |
|  | qCTS10a | 10 | 25.2 | RM239<br>(RM311) | 2,2 | Scadj |
|  | qCTS10b | 10 | 118.3 | RM333 | 2,75 | Scadj |
|  | qCTS11a | 11 | 50.8 | RM479 | 3,1 | Scadj |
|  | qCTS12a | 12 | 43.2 | RM512 | 3,7 | CH, CFL,<br>HT16adj,<br>HT23adj,<br>FL23adj |
|  | qCTS12b | 12 | 74.5 | RM309 | 2,7 | CH |

qCTS= Putative QTLs associated with cold tolerance in rice at the seedling stage, the numbers correspond to chromosomes on which they were detected and letters order. HT16adj: height adjusted before screening; FL16:leaf length adjusted before screening; HT23adj: height adjusted after screening; FL23adj: leaf length adjusted after screening, CH: growth in height; CFL: leaf growth; Scadj: adjusted score.

## DISCUSSION

Height and leaves lengths measurements before and after screening indicate that there’s been an increase in growth despite abiotic stress conditions in which seedlings were submitted. Thus, that’s allowed to speculate on presence of QTLs associated with cold tolerance in these segregating populations. Heritability values before and after stress showed higher value before stress. This difference is explained by presence of stress factor and the fact that all the seedlings put in the same stress conditions did not had the same response, which allowed after screening to classify F_2_ as sensitive, moderately tolerant and tolerant depending on leaves colors. High heritability values confirm that screening was used to determine differences between genotypes by minimizing the effect of the environment. This means, that good data quality can be used to identify QTLs associated with cold tolerance at the seedling stage. Different models obtained gave us correlations (R) value greater than 0.5, at the threshold p < 0.001. This indicates the strong link between studied parameters in these models. Two types of models have been presented: firstly, a linear relationship between height and leaf length before and after screening, with an improvement in the relationship after screening. And secondly a quadratic relationship between height and growth in height after screening, and between leaves length and growth rate in leaf length. Hence, growth in plant height and leaf length growth are closely linked.

Gene diversity index calculated allowed to have the most discriminative markers. Minimum index for a marker to be considered highly discriminative being 0.72 [21]. This brought together the varieties according to their genetic proximity. These indices were then allowed to say in this study that 13 markers would be on the loci harboring major QTLs associated with cold tolerance in rice at the seedling stage. Phylogenetic tree obtained through the study of allelic diversity of 30 rice varieties at seedling stage was used to validate the results of phenotypic studies in low temperature conditions [11]. Thus, this tree presents two major distinct groups, a large group of susceptible varieties and another of tolerant varieties.

Putative QTLs were detected some analysis was performed with program SIM and CIM, LOD ≥ 2 was fixed. Thus, for SIM program number of QTLs was 14 while; for CIM program number of QTLs was 15. This gives a difference of one QTL between the two programs performed, CIM program with a putative QTL in more at the chromosome 6 at the location of marker RM454. The QTL with the highest LOD (6.6) on chromosome 9 is closely linked to the marker RM215. Other putative QTLs with interesting value of LOD were located on chromosomes 1, 6, 11 and 12 (with respective LOD of 3.6, 3, 3.1 and 3.7), and tightly linked respectively to markers RM250, RM528, RM479 and RM512. This study allowed detecting 15 putative QTLs. Identification of QTLs responding more to CIM that allows having flanking markers for one or more QTLs, thus number of putative QTLs detected in our study was approved by the CIM program. Among these 15 QTLS detected, 3 were detected for height and leaf length before stress; 8 were detected only for parameters after stress, and 4 were detected for the parameters before and after stress. Hence, 12 putative QTLs were associated with parameters taken after screening; there are putative QTLs associated with cold tolerance at the seedling stage.

## CONCLUSION

Thus, this study confirmed association of major QTLs with cold tolerance in rice at seedling stage. These QTLs are tightly linked to the most discriminative markers RM335, RM348, RM261, RM241 and RM255 markers. Moreover, this study has validated phenotypic classification of previous studies about the susceptibility of some varieties by separating cold sensitive varieties and cold tolerant genetically. In addition, studies on segregating F_2_ population (NERICA L19 X Diamante) has detected 15 putative QTLs associated with cold tolerance in rice at the seedling stage, 3 on chromosome 1 and 6 (qCTS1a, qCTS1b, qCTS1c, and qCTS6a, qCTS6b, qCTS6c); 2 on chromosomes 10 and 12 (qCTS10a, qCTS10b, and qCTS12a, qCTS12b); 1 on chromosomes 3, 4, 7, 9 and 11 (qCTS3a, qCTS4a, qCTS7a, qCTS9a, and qCTS11a) with LOD ≥ 2. Putative QTL with the largest effect (qCTS9a) on chromosome 9 is tightly linked to marker RM215, with a LOD = 6.6. This allows conjecturing that Diamante would be a good donor for cold tolerance.

## ACKNOWLEDGMENTS

To the Sahel Regional Station of AfricaRice based in Saint Louis (Senegal) for the fellowship.

